# Hydrological Extremes and their Association with ENSO Phases in Ethiopia

**DOI:** 10.1101/681528

**Authors:** Abu Tolcha Gari

**Affiliations:** Ethiopian Institute of Agricultural Research

**Keywords:** Drought, ENSO Phase, Flood, Standard Precipitation Index (SPI)

## Abstract

Ethiopia is a rain fed agriculture country, which is subjected to high climate variability in space and time, leading to hydrological extremes causing loss of life and property more frequently. Droughts are more common and sometime floods are experienced in various parts of the country. Being a tropical country, the inter-annual climate variability in Ethiopia is dominated by ENSO (ElNino and Southern Oscillation).

In this study, an attempt has been made to determine the occurrence of droughts and floods on monthly basis, by calculating the monthly SPI (Standardized Precipitation Index) using the available rainfall data during (1975-2005) at selected 26 stations that spread across the country. Based on the monthly SPI values computed, the droughts and floods of different intensities; extreme, severe and dry have been determined for all stations. The frequencies of the droughts and floods on monthly scale during the two rainy seasons, Belg (Feb-May) and Kiremt (Jun-Sep) seasons have been determined. For instance, during Belg season, there were 11 extreme (SPI < −2.0) droughts at Nazreth, 10 severe (SPI between −1.99 and −1.50) droughts at Diredawa and 14 moderate (SPI between −1.49 and −1.00) droughts at Kulumsa. Total number of droughts of all intensities over the study period is also highest (22) at Kulumsa and lowest (8) at Mekelle. Moreover, at Kulumsa both the numbers of droughts (22) and floods (22) during Belg are more, which shows that the rainfall variability is the highest at this station.

The association of the hydrological extremes during the two rainy seasons, belg and kiremt with the ENSO phases has also been examined, which forms a basis for the prediction of the occurrence of droughts (dry conditions) and floods (wet conditions) at individual stations using ENSO phases.

## Introduction

Climate variability refers to variations in the mean state and other climate statistics (standard deviation, the occurrence of extremes like drought and floods, etc) on all temporal and spatial scales beyond those of individual weather events (Selvaraju Ramasamy and Stephan Baas, 2007). Variability may result from natural or internal processes within the climate system (internal variability) or from variations in natural or anthropogenic external forces (external variability). Variability in a normally distributed parameter occurs when changes in the mean, the variance, or both cause the probability distribution to ‘shift” resulting in changes in the frequency of occurrence of extreme events in either the upper or lower tail of the distribution.

Hydrological extremes like extreme rainfall events, floods and droughts are normal, recurring features of climate; which occur virtually in all climatic regions (Wilhite 1993). They occur in high as well as low rainfall areas. Drought is a temporary aberration, in contrast to aridity, which is a permanent feature of the climate and is restricted to low rainfall areas while flood is a permanent feature of the climate and restricted to high rainfall. Drought is the consequence of a natural reduction in the amount of precipitation received over an extended period of time, usually a season or more in length. Drought is also related to the timing (i.e., principal season of occurrence, delays in the start of the rainy season, occurrence of rains in relation to principal crop growth stages) and the effectiveness of the rains (i.e., rainfall intensity, number of rainfall events). Drought differs from other natural hazards in several ways. First, since the effects of drought often accumulate slowly over a considerable period of time and may linger for years after the termination of the event, the onset and end of drought is difficult to determine. Because of this, drought is often referred to as a creeping phenomenon (Tannehill 1947). Although Tannehill first used this terminology more than fifty years ago, climatologists continue to struggle with recognizing the onset of drought and scientists and policy makers continue to debate the basis (i.e., criteria) for declaring an end to a drought. Second, the absence of a precise and universally accepted definition of drought adds to the confusion about whether or not a drought exists and, if it does, its degree of severity.

Similarly, floods are the most common phenomenon that causes human suffering, inconvenience and widespread damage to buildings, structures, crops and infrastructures. Floods have been observed to disrupt personal, economic & social activities and set back a nations security & development by destroying roads, buildings and other assets. The recent major recorded flood disaster that still lingers in our mind is the (2006) flood in Ethiopia. During 2006 flood in Ethiopia, the flooding occurred in almost all parts of the country. In the North, localities in Tigray and in the northeast, Amhara region have been affected by emerging floods. In the south and East, the major flood damage was registered with loss of huge number of human and animal lives, loss of property. In the South, the Baro River was swelling to create a flood situation (Semu, 2007).

Droughts and floods are extreme hydrological events that may adversely affect the social, economic, cultural, political and other functions of a region. Drought and flood predictions may prevent these adverse consequences to a significant extent. In order to reach such a target, it is necessary to develop a method of prediction based on the available past experiences as well as on environmental conditions.

The Horn of Africa is known for frequent and severe droughts. On average, a severe drought may be anticipated once every couple of years during the months that comprise the season (s). The impact of hydrological extremes like drought and flood on society and agriculture is a real issue but it is not easily quantified. Reliable indices to detect the spatial and temporal dimensions of flood and drought occurrences and their intensity are necessary to assess the impact and also for decision-making and crop research priorities for alleviation (Seiler et al., 1998).

ENSO was found to be one of the main factors that lead to climate variability in Ethiopia (Bekele F. 21997; Woldegiorgis *et al*., 2000). Many researchers now believe that the occurrence of various droughts in Africa, especially in Southern Africa and the Horn, are caused by physical processes related to the occurrence of ENSO events thousands of miles away. If valid and reliable information about the linkages between these occurrences becomes available, it could help to forecast Sub-Saharan African droughts. Empirical data indicate an association between ENSO events and droughts in Ethiopia. Thus, an ENSO-based early warning system, used effectively by policymakers, could help to reduce the societal impacts of droughts as well as floods in Ethiopia.

It have been suggested that monitoring the Southern Oscillation Index (SOI), which is the atmospheric component of ENSO phenomenon, could predict the hydrological extremes with a longer lead time. It has been observed (Stone et al, 1996) that the SOI phase as determined by the change in average monthly SOI over the two previous months can give future seasonal rainfall probabilities more accurately than using SOI averages. Therefore, this study aims at determining the frequencies of droughts and floods at various rainfall stations in Ethiopia using the Standardized Precipitation Index (SPI) and examining their association with SOI phases within the historical record of rainfall data.

## Materials and Methods

### Data used in the study

Monthly rainfall datasets have been acquired from the National Meteorological Agency (NMA) for a period of 31 years during 1975-2005 for 26 stations spread across the country (Fig. 1). Monthly rainfall for all these rain stations has been used to derive Standardized Precipitation Index (SPI). For the other 5 stations, as the data is missed continuously for 3 or more years, they have been omitted in the study.

**Figure 1.**
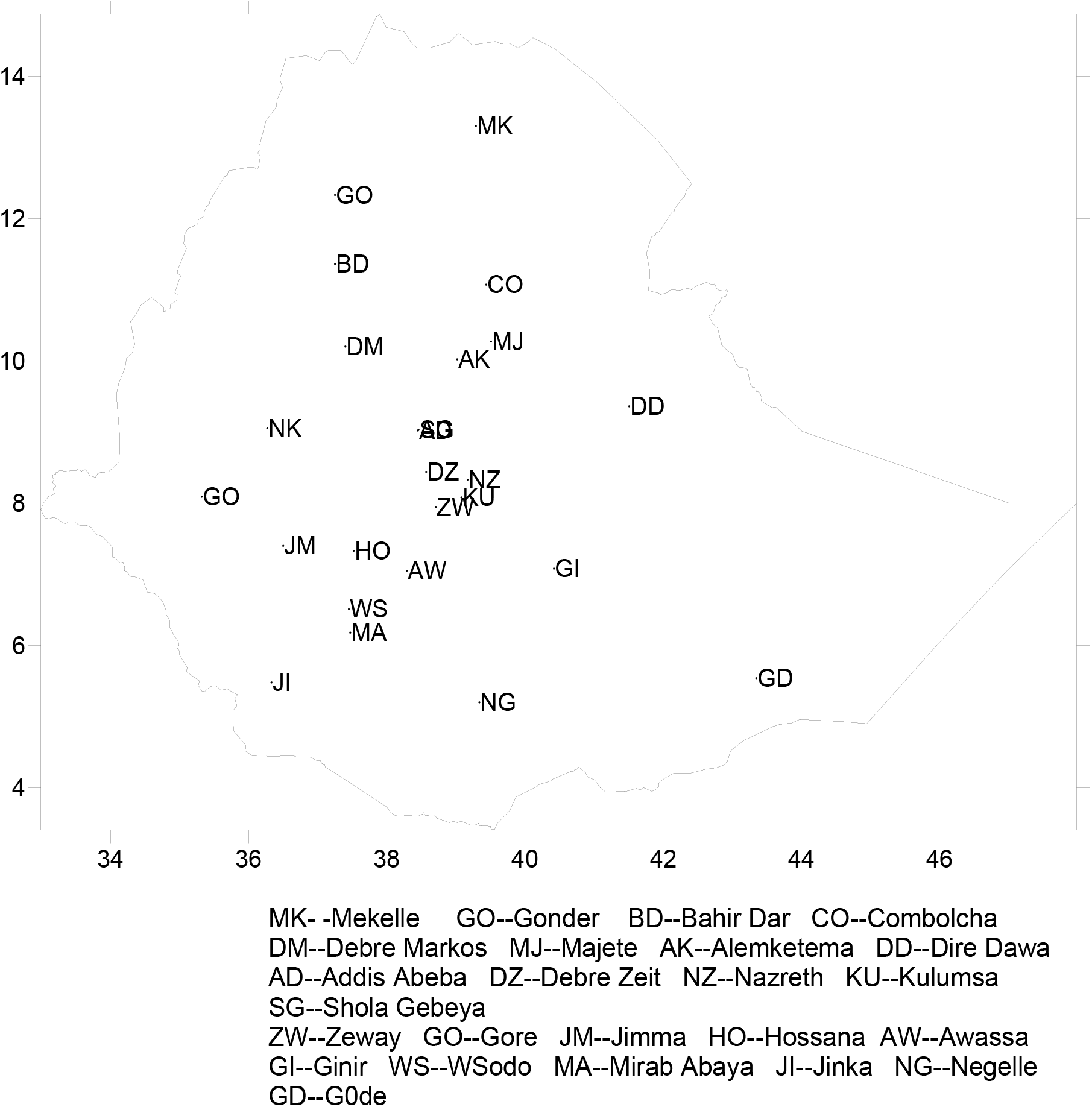
Distribution of Rainfall Stations

For a few of the selected 26 stations, the missing monthly rainfall data (which is around 2%) has been supplemented with the corresponding mean monthly rainfall of that station. Similarly, the monthly SOI phases during the period 1975-2005 have been collected from the Long Paddock web page (http://www.longpaddock.qld.gov.au/Help/SOIPhases/index.html).

## Methods

### Standard Precipitation Index

The SPI was chosen for this study because of its simplicity and being based solely on the available precipitation data. The SPI method was first developed by Mckee et al. (1993) transforms the precipitation parameter to a single numerical value for defining the drought and flood condition of areas with different climates. The SPI allows the determination of duration, magnitude and intensity of droughts and floods (Hayes et al., 1999). Its main advantage is that it can be calculated for several time scales (McKee et al., 1995) and identifies various drought types: hydrological, agricultural or environmental. This index enjoys several advantages over the others. The nature of the SPI allows an analyst to determine the rarity of a drought or an anomalously wet event at a particular time scale for any location in the world that has a precipitation record.

Therefore, SPI is calculated from monthly precipitation record by first fitting the gamma probability distribution function and then transforming into a normal distribution so that the mean SPI is set to zero (McKee et al., 1993; Edwards and McKee, 1997). Positive and negative SPI values indicate wet and dry conditions, respectively. The alpha and beta parameters of the gamma probability density function are estimated for each station, for each time scale of interest (1, 3, 6, 9, 12 months, etc.), and for each month of the year. The gamma distribution is defined by its frequency or probability density function:

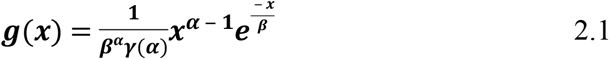

Where α and β are the shape and scale parameters respectively, x is the precipitation amount and ***γ*** (*α*) is the gamma function. Maximum likelihood solutions are used to optimally estimate α and β:

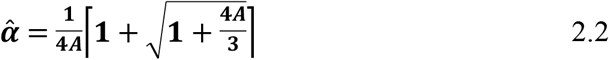

Where:

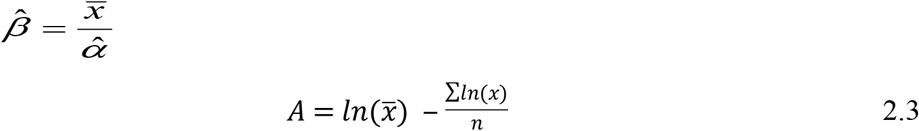

*Where n* = number of precipitation observations

The resulting parameters are then used to find the cumulative probability of an observed precipitation event for the given month and time scale for the station in question. The cumulative probability is given by:

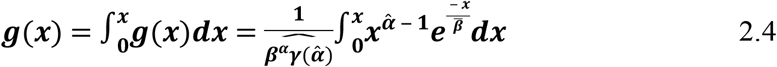

Letting 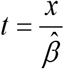, this equation becomes the incomplete gamma function:

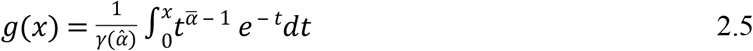

Since g(x) is undefined for x=0 and a precipitation distribution may contain zeros, the cumulative probability becomes:

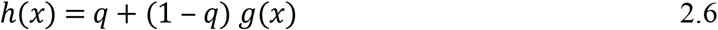

Where q is the probability of a zero and g(x) is the cumulative probability of the incomplete gamma function. If m is the number of zeros in a precipitation time series, then q can be estimated by m/n. By applying Eq. (2.6), errors are eventually introduced to parameters α and β of gamma distribution. These errors depend on the number of months with null precipitation (x=0) and they are evident only for the 1-month precipitation. For larger time scales (e.g. 3-month, 6-month, etc.) the probability of null precipitation was zero.

The cumulative probability, h(x), after its computation, is transformed to the standard normal random variable z with mean equal to zero and variance of one, which is the value of the SPI. Once standardized the strength of the anomaly is classified as set out in Table 1. This table also contains the corresponding probabilities of occurrence of each severity arising naturally from the normal probability density function. Thus, at a given location for an individual month, moderate dry periods (SPI <=−1) have an occurrence probability of 9.2%, whereas extreme dry periods (SPI<=−2) have an event probability of 2.3%. Extreme values in the SPI will, by definition, occur with the same frequency at all locations.

**Table 1.** Classification by SPI values and corresponding event probabilities (%)

| SPI value | Category | Probability (%) |
| --- | --- | --- |
| 2.00 or more | Extremely wet | 2.3 |
| 1.5 to 1.99 | Severely wet | 4.4 |
| 1.00 to 1.49 | Moderately wet | 9.2 |
| -0.99 to 0.99 | Near normal | 68.2 |
| -1.49 to -1.00 | Moderately dry | 9.2 |
| -1.99 to -1.50 | Severely dry | 4.4 |
| -2 or less | Extremely dry | 2.3 |

An analyst with a time series of monthly precipitation data for a location can calculate the SPI for any month in the record for the previous *i* months where *i*=1, 2, 3, 12, 24, 48, depending upon the time scale of interest. Hence, the SPI can be computed for an observation of a 3 month total of precipitation as well as a 48 month total of precipitation. For this study, a 3 month and 6 month SPI is used for a short-term or seasonal drought index, a 12 month SPI is used for an intermediate-term drought index. Therefore, the SPI for a month/year in the period of record is dependent upon the time scale.

## Results and Discussion

Based on the monthly SPI values computed over the period 1975-2005, the droughts (dry conditions) and floods (wet conditions) of different intensities; extreme, severe and dry (Table-1) was determined for the 26 stations considered in the study.

### Droughts

The drought frequencies during Belg, Kiremt and Bega over the study period have been presented in graphs (Fig.2).

**Figure 2.**
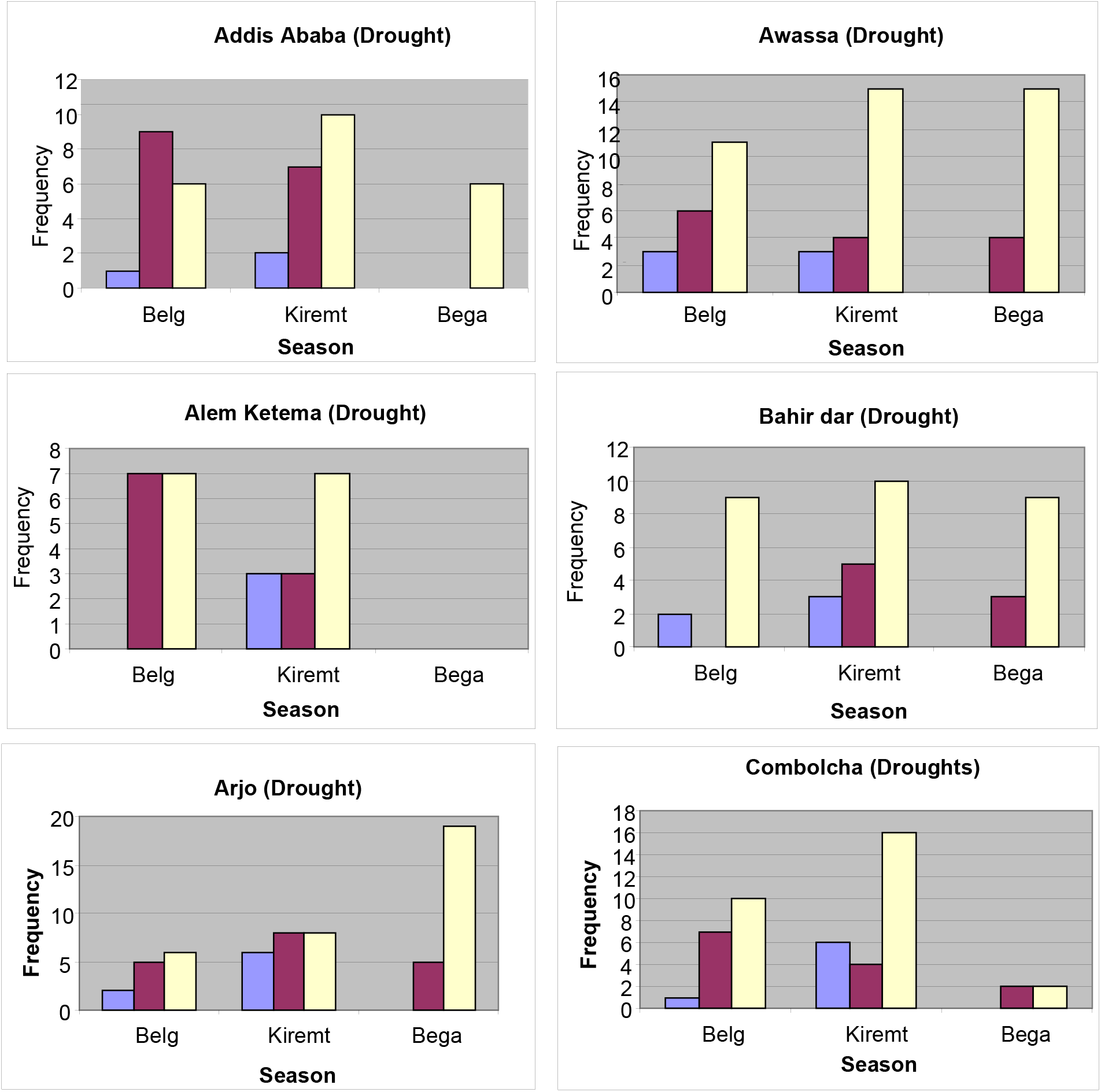
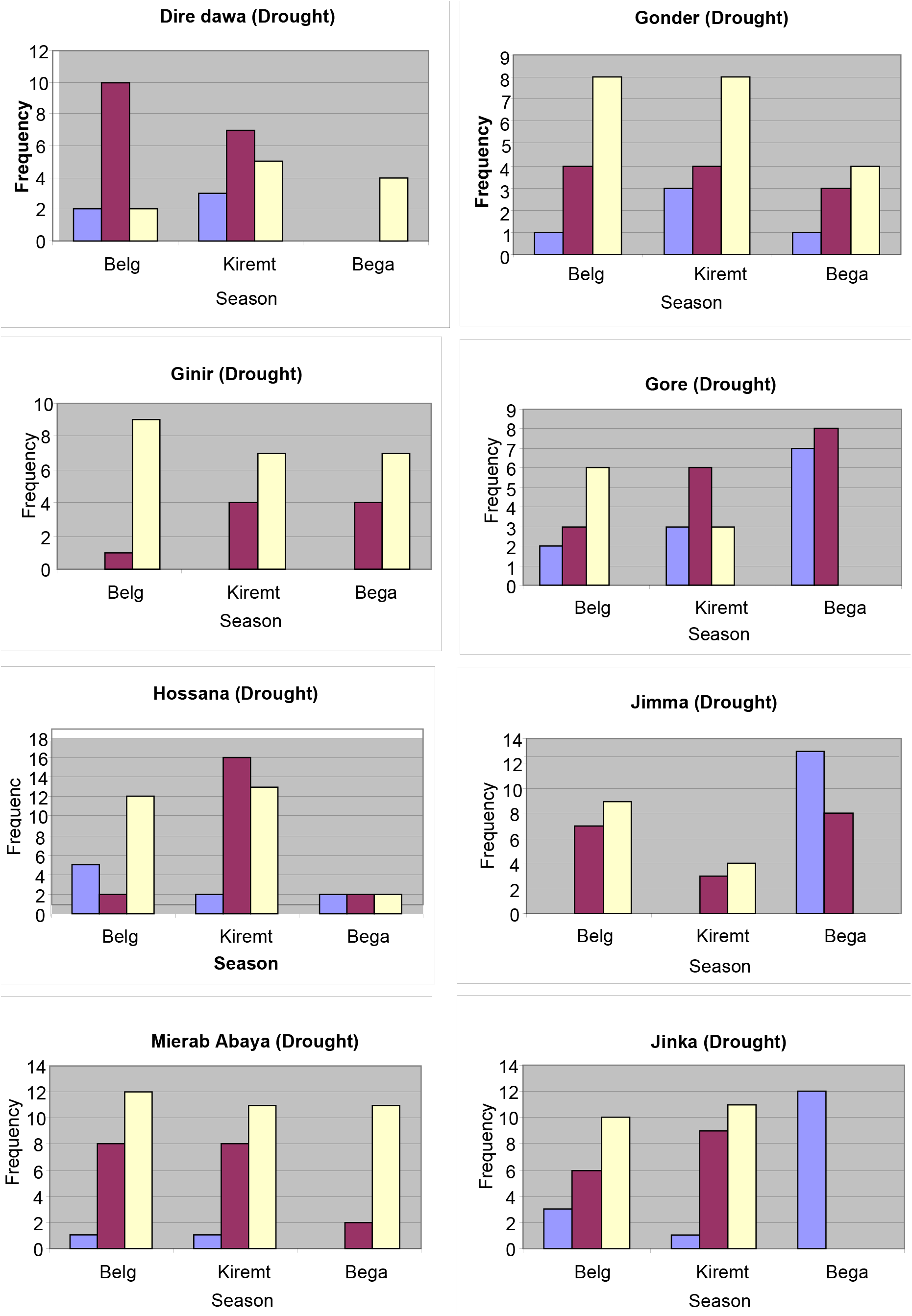
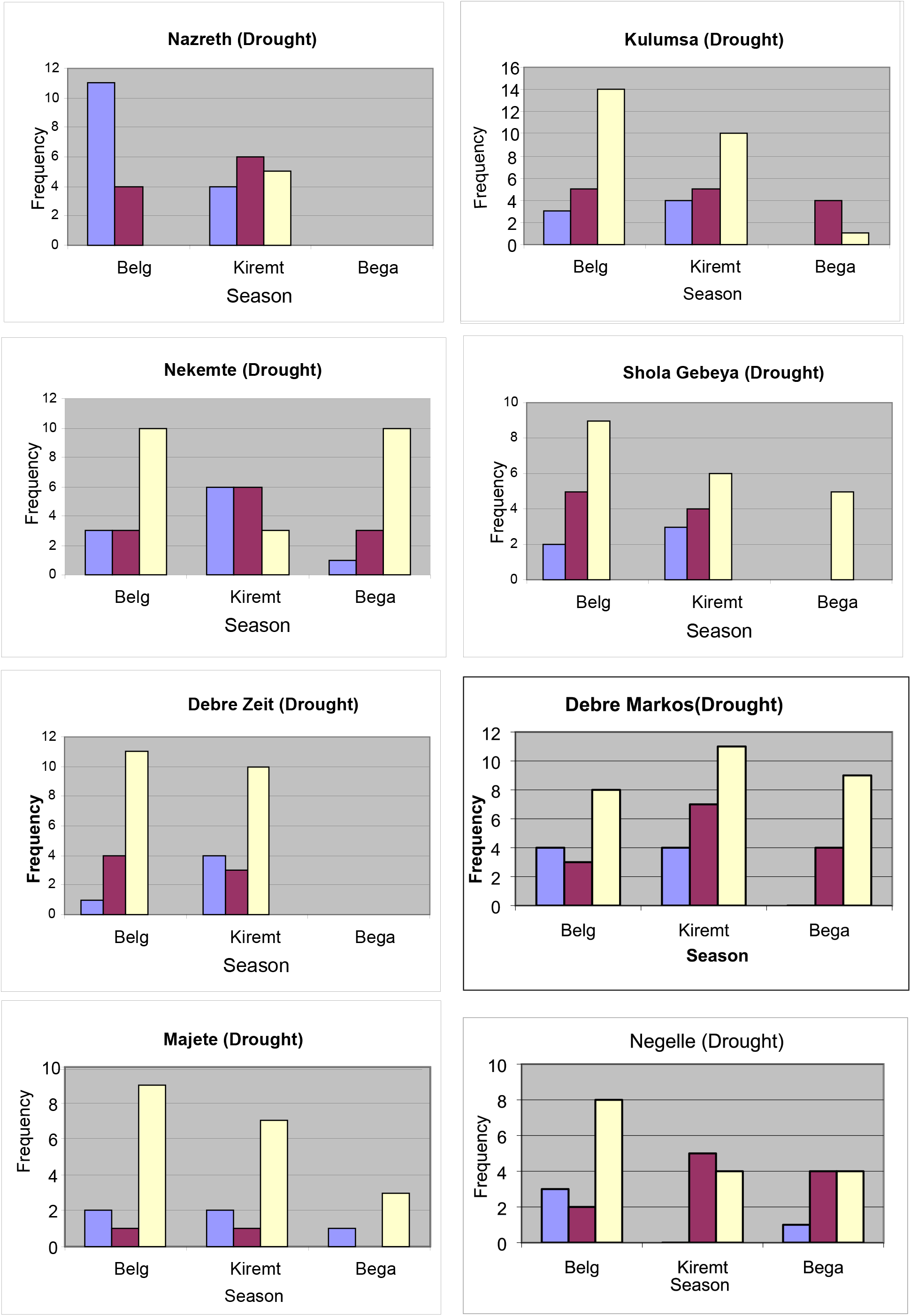
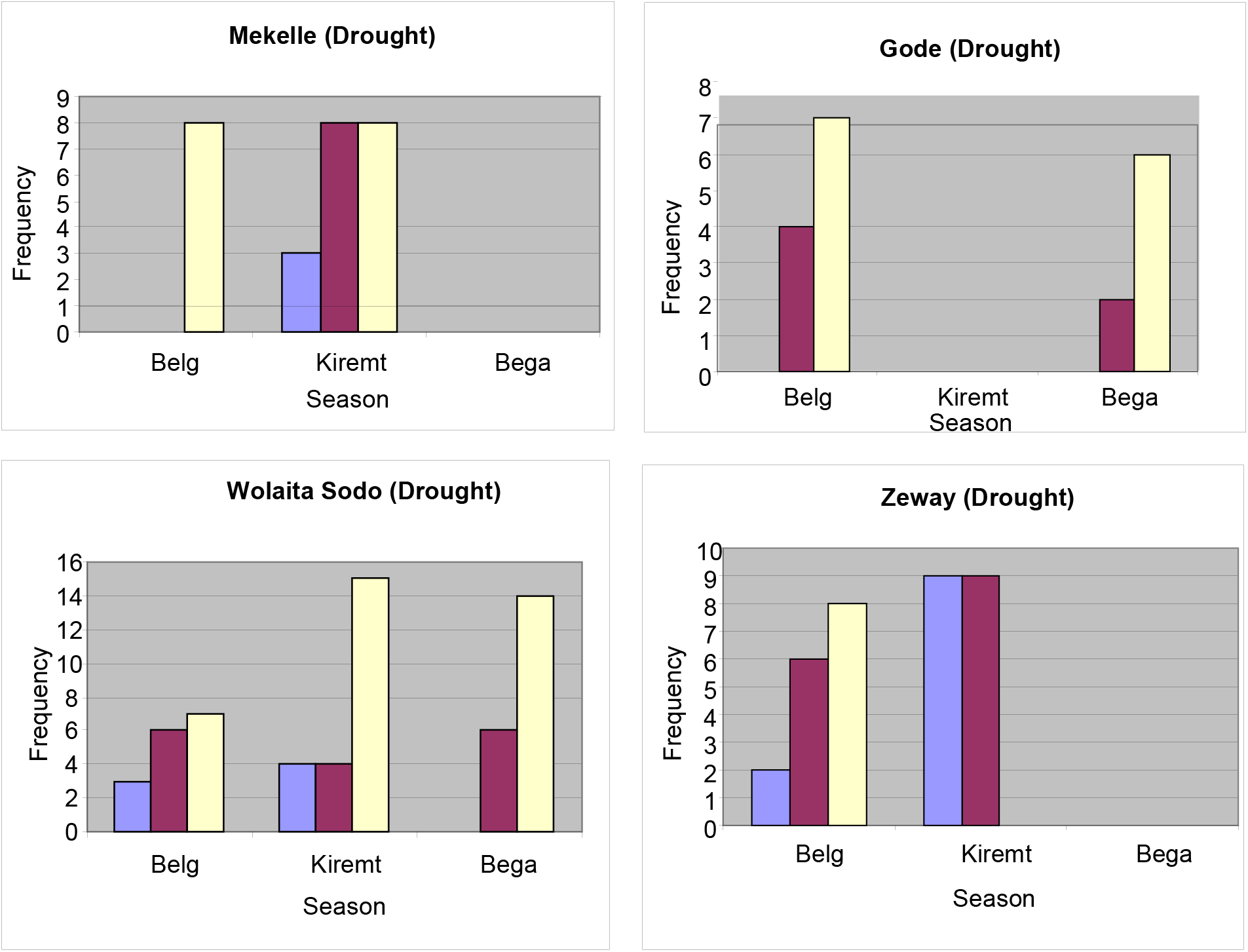
Frequencies of Droughts of different intensities (Extreme – Blue; Severe – Brown and Moderate – Yellow) over the period 1975-2005

Figure-2 shows that, during belg season, there were 11 extreme (SPI < −2.0) droughts at Nazreth, 10 severe (SPI between −1.99 and −1.50) droughts at Diredawa and 14 moderate (SPI between −1.49 and −1.00) droughts at Kulumsa. Total number of droughts of all intensities over the period 1975-2005 is also highest (22) at Kulumsa and lowest (8) at Mekelle. It can also be observed from the figure that, during belg extreme drought was absent at 5 stations, severe drought was absent at 2 stations and moderate drought was absent at Mekelle.

Similarly during kiremt season, Zeway experienced 9 extreme droughts, while Hosana experienced 16 severe and Combolcha experienced 16 moderate droughts over the period 1975-2005. The total number of droughts during Kiremt is highest (31) at Hosana. During this season, extreme droughts were absent at 4 stations, severe droughts were absent at Gode and moderate droughts were absent at Gode and Zeway. Surprisingly, there were no droughts at all at Gode during kiremt over the period of study.

### Floods

The flood frequencies (of different intensities) during belg, kiremt and bega over the study period have been presented in graphs (Fig. 3).

**Fig. 3.**
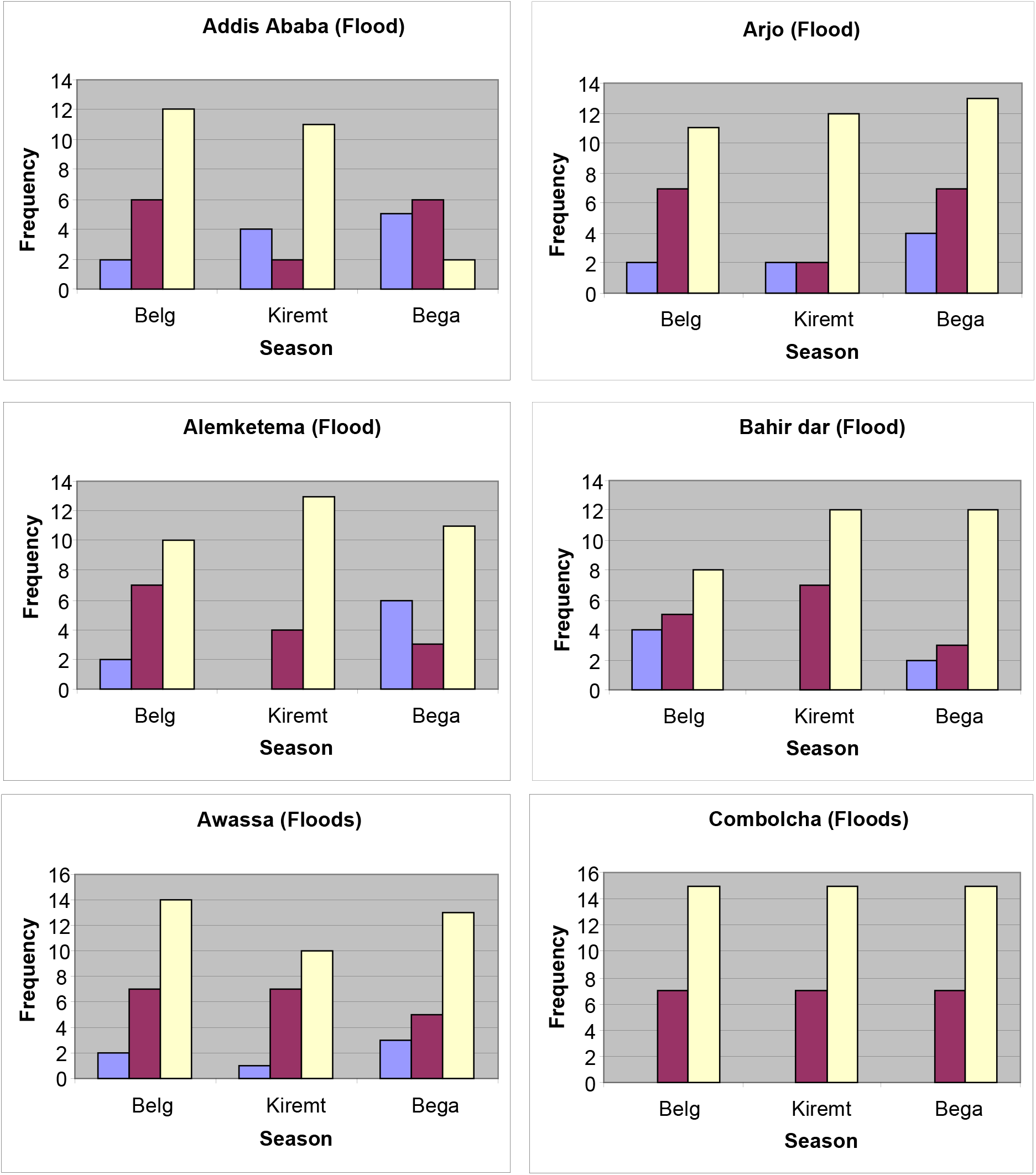
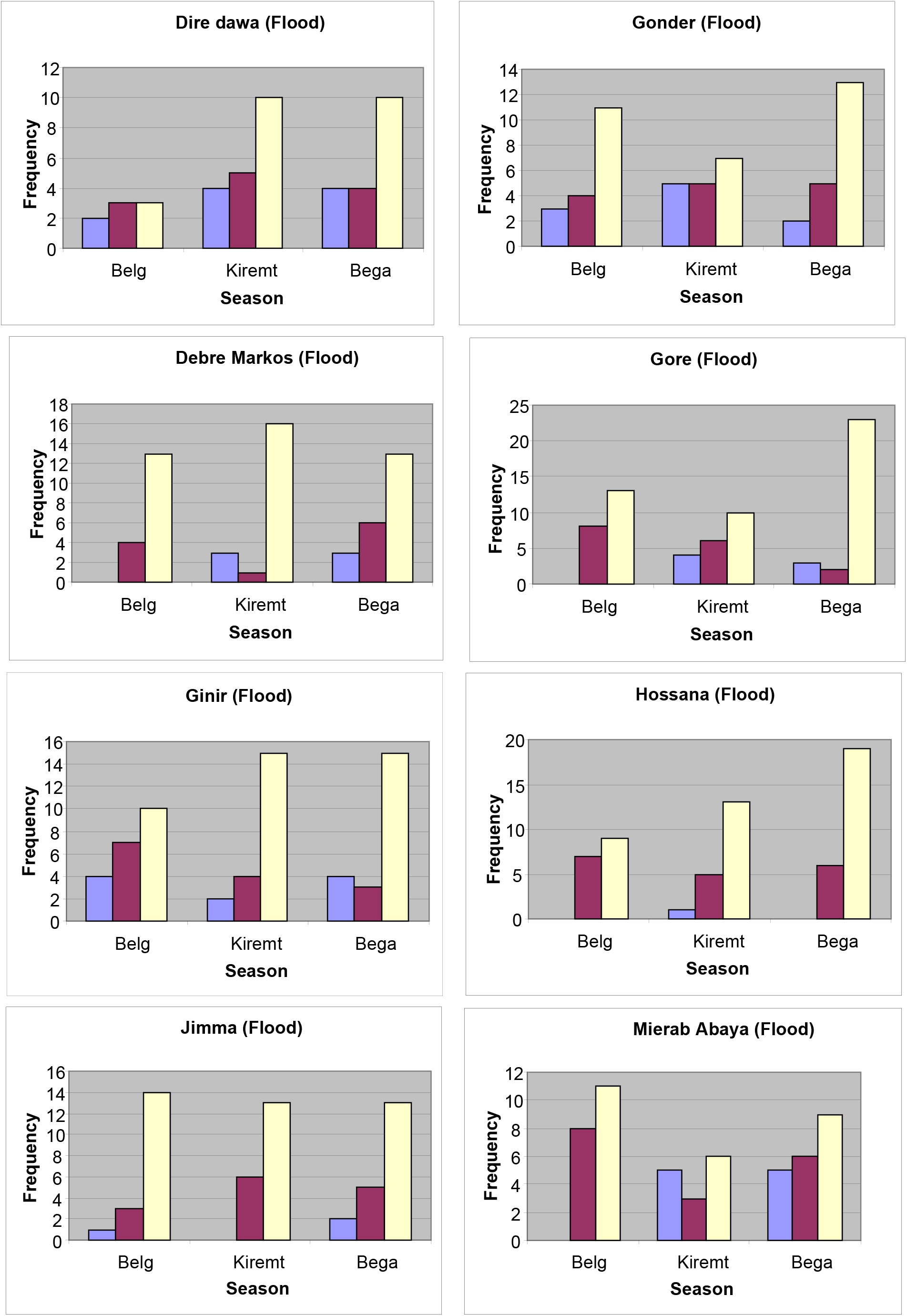
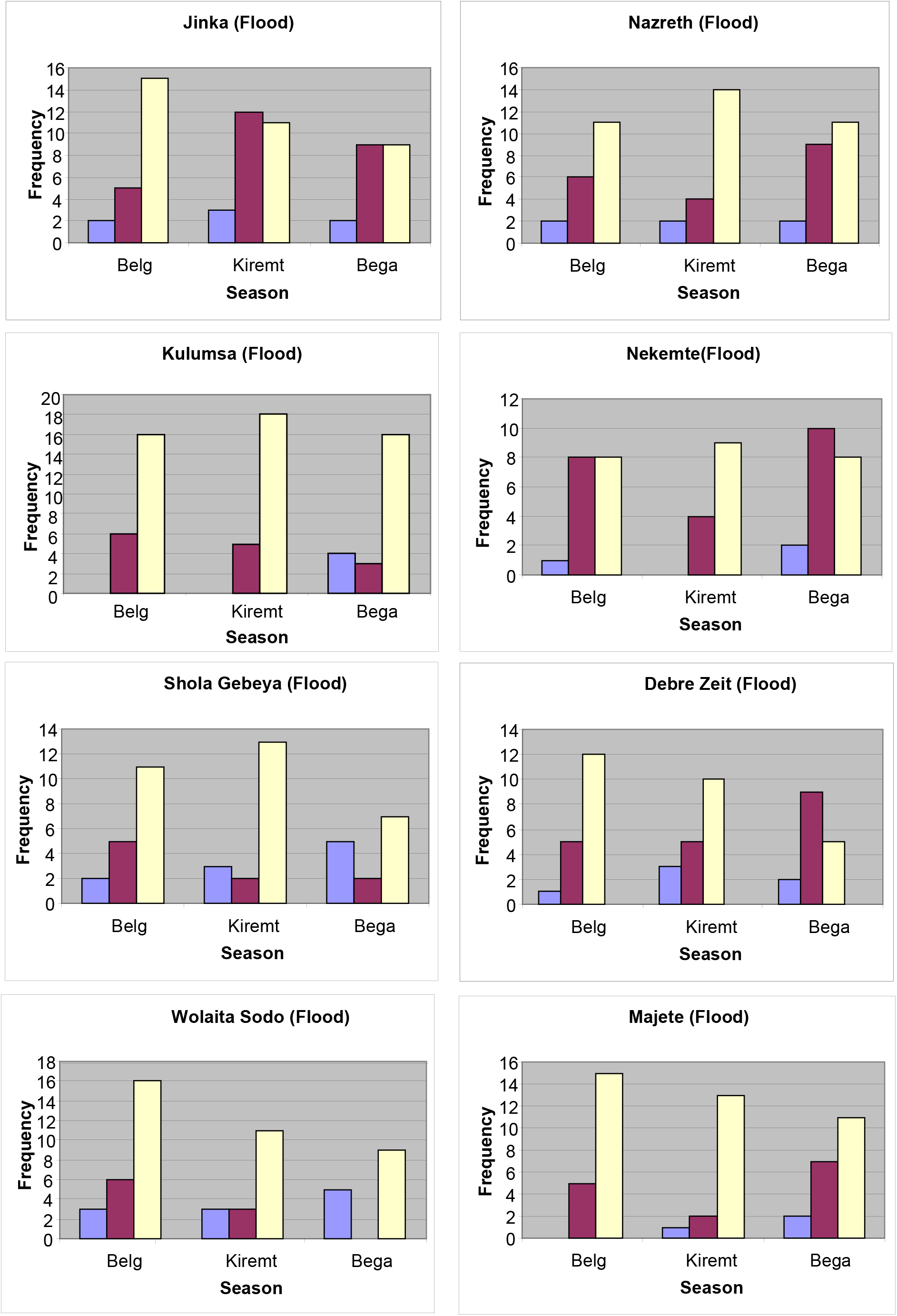
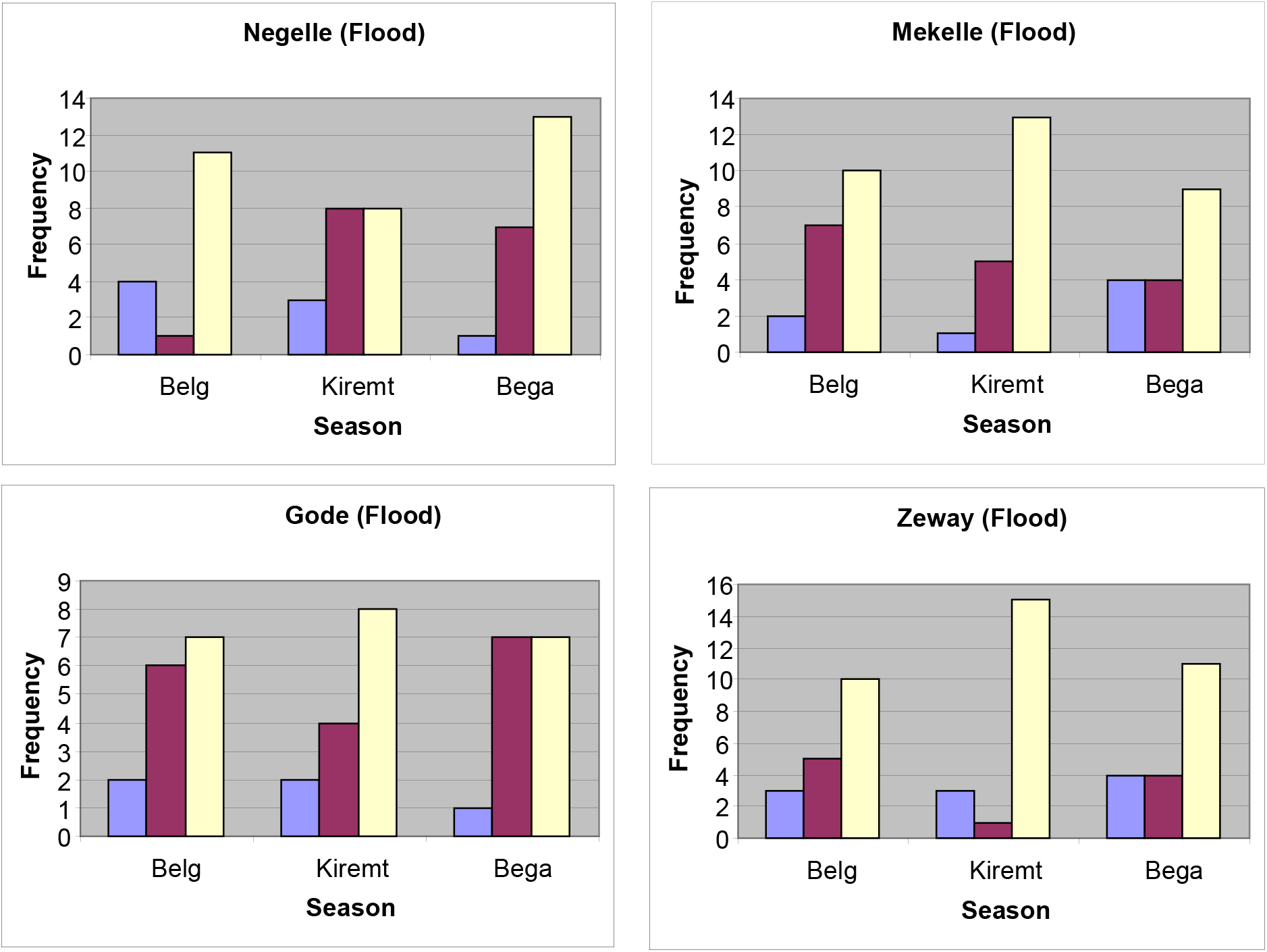
Frequencies of Floods of different intensities (Extreme – Blue; Severe – Brown and Moderate – Yellow) over the period 1975-2005

As shown in figure-3, during belg season, 3 stations, Bahirdar, Ginir and Negelle have experienced the highest number (4) of extreme (SPI > 2.0) floods, while Gore, Mirab Abaya and Nekemte have experienced 8 severe (SPI between 1.50 and 1.99) floods each and Kulumsa and Wolaita Sodo have experienced 16 moderate (SPI between 1.00 and 1.49) floods. The total number of floods of all intensities during belg over the study period is the highest (25) at Wolaita Sodo and lowest (8) at Diredawa (Fig.-3). Thus, at Kulumsa both the numbers of droughts (22) and floods (22) during belg are more, which shows that the rainfall variability is the highest at this station.

Similarly, during kiremt, Gonder and Mirab Abaya experienced the highest number (5) of extreme floods, while Jinka experienced 12 severe floods and Kulumsa experienced 18 moderate floods over the period. The total number of floods of all intensities is highest (26) at Jinka and lowest (8) at Gode. Thus, at Gode both the numbers of droughts (0) and floods (8) are the lowest, during kiremt, which shows that the rainfall variability is the lowest at this station. Only extreme floods were absent at 6 stations, while all the stations have experienced the severe and moderate floods during this season.

### Hydrological Extremes and Their Association with ENSO

In this section, at each of the 26 stations considered in the study, the association of the hydrological extremes, namely the droughts and floods with ENSO has been examined on monthly basis. For this, the monthly occurrences of the hydrological extremes during the two rainy seasons, belg (Feb-May) and kiremt (Jun-Sep) associated with each of the five SOI phases (namely 1, 2, 3, 4 and 5) for all the months (starting from January) preceding and up to the last month have been evaluated over the period 1975-2005. As an example, these frequencies for Addis Ababa have been presented in Table-2.

**Table 2.**
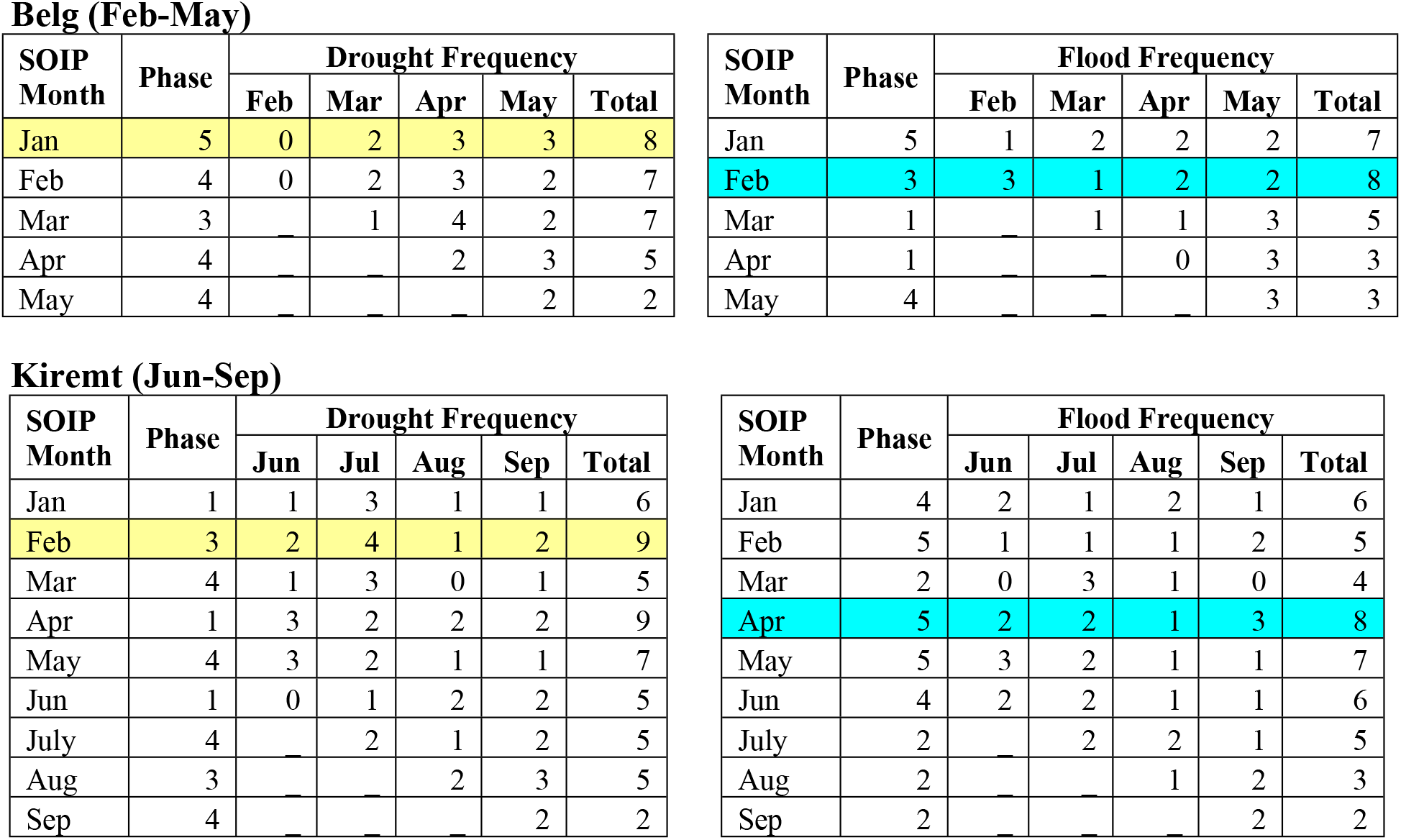
Drought and Flood Frequencies associated with SOI phases during Belg and Kiremt for Addis Ababa

Belg (Feb-May)
| SOIP Month | Phase | Drought Frequency |  |  |  |  |
| --- | --- | --- | --- | --- | --- | --- |
|  |  | Feb | Mar | Apr | May | Total |
| Jan | 5 | 0 | 2 | 3 | 3 | 8 |
| Feb | 4 | 0 | 2 | 3 | 2 | 7 |
| Mar | 3 | — | 1 | 4 | 2 | 7 |
| Apr | 4 | — | — | 2 | 3 | 5 |
| May | 4 | — | — | — | 2 | 2 |

| SOIP Month | Phase | Flood Frequency |  |  |  |  |
| --- | --- | --- | --- | --- | --- | --- |
|  |  | Feb | Mar | Apr | May | Total |
| Jan | 5 | 1 | 2 | 2 | 2 | 7 |
| Feb | 3 | 3 | 1 | 2 | 2 | 8 |
| Mar | 1 | — | 1 | 1 | 3 | 5 |
| Apr | 1 | — | — | 0 | 3 | 3 |
| May | 4 | — | — | — | 3 | 3 |

Kiremt (Jun-Sep)
| SOIP Month | Phase | Drought Frequency |  |  |  |  |
| --- | --- | --- | --- | --- | --- | --- |
|  |  | Jun | Jul | Aug | Sep | Total |
| Jan | 1 | 1 | 3 | 1 | 1 | 6 |
| Feb | 3 | 2 | 4 | 1 | 2 | 9 |
| Mar | 4 | 1 | 3 | 0 | 1 | 5 |
| Apr | 1 | 3 | 2 | 2 | 2 | 9 |
| May | 4 | 3 | 2 | 1 | 1 | 7 |
| Jun | 1 | 0 | 1 | 2 | 2 | 5 |
| July | 4 | — | 2 | 1 | 2 | 5 |
| Aug | 3 | — | — | 2 | 3 | 5 |
| Sep | 4 | — | — | — | 2 | 2 |

| SOIP Month | Phase | Flood Frequency |  |  |  |  |
| --- | --- | --- | --- | --- | --- | --- |
|  |  | Jun | Jul | Aug | Sep | Total |
| Jan | 4 | 2 | 1 | 2 | 1 | 6 |
| Feb | 5 | 1 | 1 | 1 | 2 | 5 |
| Mar | 2 | 0 | 3 | 1 | 0 | 4 |
| Apr | 5 | 2 | 2 | 1 | 3 | 8 |
| May | 5 | 3 | 2 | 1 | 1 | 7 |
| Jun | 4 | 2 | 2 | 1 | 1 | 6 |
| July | 2 | — | 2 | 2 | 1 | 5 |
| Aug | 2 | — | — | 1 | 2 | 3 |
| Sep | 2 | — | — | — | 2 | 2 |

As shown in the table, when the SOI is consistently near zero (phase-5) in January there were 8 drought cases (of all intensities) during Belg season at Addis Ababa, over the study period. Similarly, when the SOI is falling (phase-3) in February, there were 8 flood cases during Belg and 9 drought cases during Kiremt. This opposite variation of ENSO with Ethiopian rainfall between belg and kiremt has been observed by several other researchers also. The table also shows that when the SOI is in phase-5 in April, there were 8 flood cases during kiremt.

Similar analysis has been carried out for the remaining 25 stations and the SOI phases associated with the highest frequency of droughts and floods at all the 26 stations considered in the study, have been summarized and presented in Appendix-I for belg and kiremt seasons.

### ENSO and Hydrological Extremes during Belg

The monthly occurrences of the hydrological extremes during belg (Feb-May) associated with each of the five SOI phases (namely 1, 2, 3, 4 and 5) for all the months (starting from January) preceding and up to the current month have been evaluated over the period 1975-2005 for the stations considered in the study. The SOI phases associated with the highest frequency of Droughts and Floods during this season are summarized in Table-3.

**Table 3.**
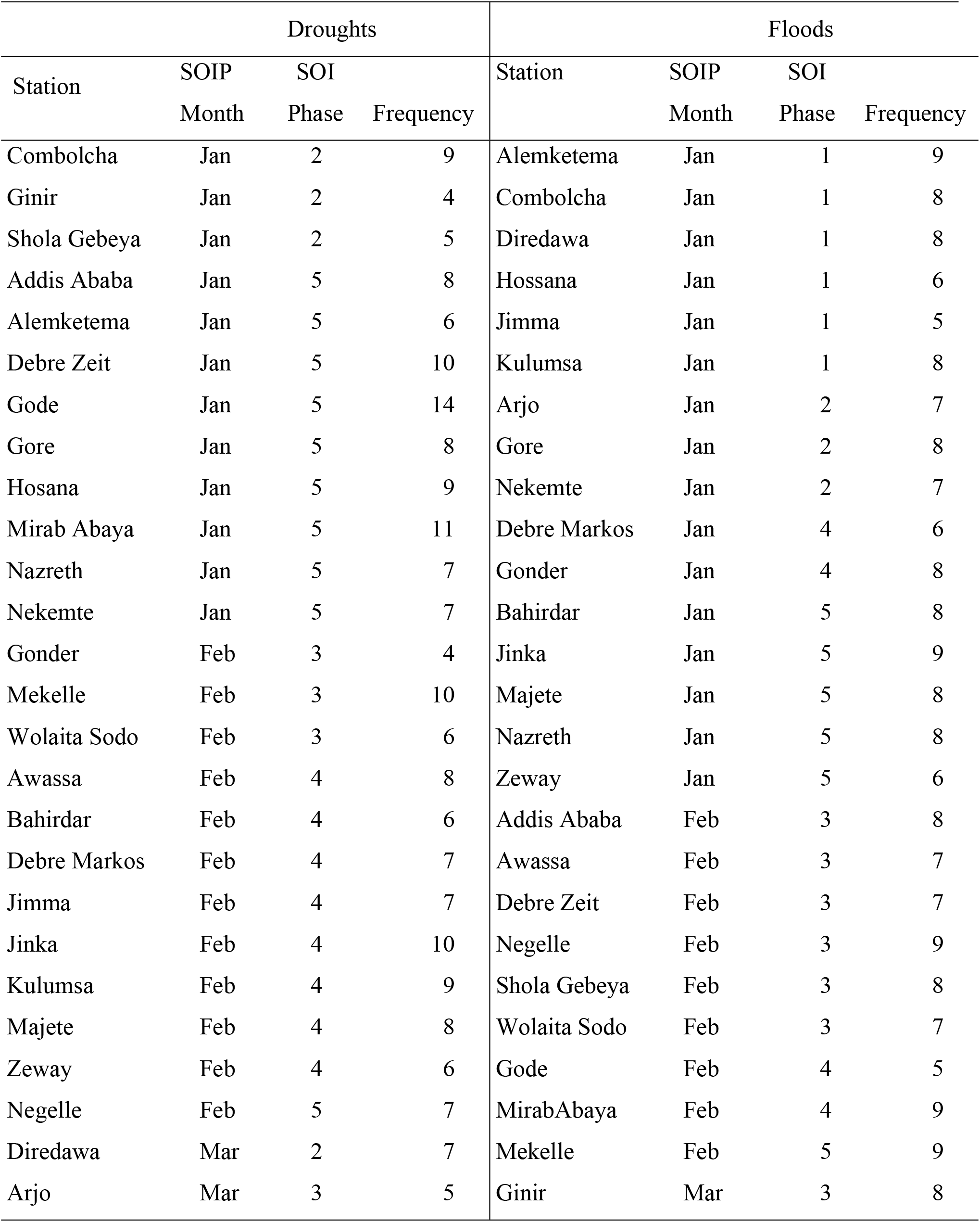
SOI phases associated with the highest frequency of Droughts and Floods at 26 stations considered in the study during Belg (Feb-May) season

As shown in the table, during belg more stations experienced the higher drought frequencies when the SOI is in phase-5 (nine stations) in January and also when the SOI is in phase-4 (eight stations) in February. Similarly, when the SOI is in phase-1 (six stations) and phase-5 (five stations) in January and in phase-3 (six stations) in February, more stations experienced the higher flood frequencies. It can be observed that when the SOI is in phase-5 in January, while nine stations experienced higher drought frequencies, six other stations have experienced higher flood frequencies. Similarly, when the SOI is in phase-5, both the drought and flood frequencies are higher at Nazreth.

### ENSO and Hydrological Extremes during Kiremt

Similarly, the frequencies of both hydrological extremes (drought and flood) during kiremt (Jun-Sep) associated with each of the five SOI phases (namely 1, 2, 3, 4 and 5) for all the months (starting from January) preceding and up to the current month have been evaluated for all stations considered in the study over the study period have been presented in Appendix-I. The SOI phases associated with the highest frequency of droughts and floods during this season are summarized in Table 4.

**Table 4.** SOI phases associated with the highest frequency of droughts and floods at 26 stations considered in the study during Kiremt (Jun-Sep) season

| Droughts |  |  |  | Floods |  |  |  |
| --- | --- | --- | --- | --- | --- | --- | --- |
| Station Name | SOIP Month | SOI Phase | Frequency | Station Name | SOIP Month | SOI Phase | Frequency |
| Jimma | Jan | 4 | 7 | Gonder | Jan | 2 | 10 |
| Wolaita Sodo | Jan | 4 | 8 | Diredawa | Jan | 4 | 8 |
| Gonder | Jan | 5 | 9 | Combolcha | Jan | 5 | 9 |
| Shola Gebeya | Jan | 5 | 7 | Debre Markos | Jan | 5 | 11 |
| Bahirdar | Feb | 2 | 9 | Mekelle | Jan | 5 | 10 |
| Addis Ababa | Feb | 3 | 9 | Nazreth | Jan | 5 | 12 |
| Combolcha | Feb | 3 | 8 | Shola Gebeya | Jan | 5 | 7 |
| Diredawa | Feb | 3 | 8 | Wolaita Sodo | Jan | 5 | 10 |
| Awassa | Mar | 1 | 10 | Bahirdar | Feb | 3 | 7 |
| Ginir | Mar | 3 | 8 | Gode | Feb | 4 | 8 |
| Hosana | Mar | 3 | 9 | Debre Zeit | Mar | 2 | 7 |
| Jinka | Mar | 3 | 13 | Kulumsa | Mar | 2 | 10 |
| Kulumsa | Mar | 3 | 9 | Mirab Abaya | Mar | 2 | 8 |
| Nekemte | Mar | 3 | 8 | Addis Ababa | Apr | 5 | 8 |
| Alemketema | Apr | 1 | 9 | Awassa | Apr | 2 | 8 |
| Debre Markos | Apr | 1 | 12 | Ginir | Apr | 3 | 8 |
| Debre Zeit | Apr | 1 | 14 | Gore | Apr | 5 | 9 |
| Arjo | May | 1 | 13 | Hossana | Apr | 1 | 9 |
| Negelle | May | 1 | 7 | Jinka | Apr | 5 | 9 |
| Zeway | May | 1 | 9 | Majete | Apr | 5 | 5 |
| Gode | May | 4 | 16 | Nekemte | Apr | 3 | 5 |
| Mekelle | May | 4 | 15 | Alemketema | May | 5 | 6 |
| Mirab Abaya | May | 4 | 11 | Negele | May | 4 | 11 |
| Nazreth | May | 4 | 9 | Arjo | Jun | 3 | 10 |
| Gore | Jun | 3 | 7 | Jimma | Jun | 3 | 13 |
| Majete | Jun | 3 | 12 | Zeway | Jun | 2 | 8 |

When the SOI is in phase-3 in March (Table-4) during kiremt season, five stations experienced the higher drought frequencies, while six stations have experienced the higher flood frequencies when the SOI is in phase-5 in January. Thus, when the SOI is in phase-5 in January, nine stations (Addis Ababa, Alemketema, Debrizeit, Gode, Gore, Hosana, MirabAbaya, Nazreth and Nekemte) have experienced droughts during Belg (Table-3), six stations (Combolcha, Debre Markos, Mekelle, Nazreth, Shola Gebeya and Wolaita Sodo) experienced floods during Kiremt (Table-4). This shows that SOI phase-5 is associated with droughts during Belg and floods during Kiremt at majority of the stations in Ethiopia. This preliminary analysis forms a basis for the prediction of hydrological extremes in Ethiopia based on the SOI phases.

## Summary and Conclusions

In this study, the frequencies of droughts and floods at 26 rainfall stations spread across Ethiopia have been determined using the Standardized Precipitation Index (SPI), which is simple and is based solely on the accessible precipitation data. Further, the association of the droughts and floods with SOI phases has been examined within the historical record of rainfall data.

Based on the monthly Standardized Precipitation Index (SPI) values computed over the period 1975-2005, the droughts (dry conditions) and floods (wet conditions) of different intensities; extreme, severe and moderate have been determined on monthly basis for all stations. It has been observed that the total number of droughts of all intensities during belg is highest (22) at Kulumsa and lowest (8) at Mekelle and during kiremt; the number of droughts is highest (31) at Hosana and nil at Gode. Similarly, the total number of floods of all intensities during belg over the study period is the highest (25) at Wolaita Sodo and lowest (8) at Diredawa and during kiremt; the number of floods is highest (26) at Jinka and lowest (8) at Gode. At Kulumsa both the numbers of droughts (22) and floods (22) during belg are more, which shows that the rainfall variability is the highest at this station. Conversely, at Gode both the numbers of droughts (0) and floods (8) are the lowest during kiremt, which shows that the rainfall variability is the lowest at this station during this season. This seasonal and spatial (at various stations) analysis of meteorological droughts and floods, provide a framework for sustainable drought monitoring and management in Ethiopia.

The association of the monthly occurrences of the hydrological extremes during the two rainy seasons, belg (Feb-May) and kiremt (Jun-Sep) with each of the five SOI phases (namely 1, 2, 3, 4 and 5) have been evaluated over the study period. Analysis of the SOI phases associated with the highest frequency of droughts and floods, has shown that during belg more stations experienced the higher drought frequencies when the SOI is in phase-5 (nine stations) in January and also when the SOI is in phase-4 (eight stations) in February. Similarly, when the SOI is in phase-1 (six stations) and 5 (five stations) in January and in phase-3 (six stations) in February, more stations experienced the higher flood frequencies. During kiremt, when the SOI is in phase-3 in February (Table-4), five stations experienced the higher drought frequencies, while six stations have experienced the higher flood frequencies when the SOI is in phase-5 in January. The above analysis clearly shows that SOI phase-5 is associated with droughts during Belg and floods during Kiremt at majority of the stations in Ethiopia. However, no spatial coherence (zone-wise) in either the frequency of the hydrological extremes or their association with the ENSO phases has been observed with this limited number of rainfall stations.

This preliminary analysis forms a basis for the prediction of hydrological extremes in Ethiopia based on the SOI phases. However, by using still larger databases, better relationships of SOI phases with the hydrological extremes can be arrived. The spatial variation and coherence of the hydrological extremes and their association with the SOI phases are also to be examined using larger databases for better prediction.

To compensate the scarcity and un-availability of long term rainfall data, remote sensing data obtained from satellites such as NDVI, can be used to compute indices based on such data like Standard Vegetation Index (SVI) or Vegetation Condition Index (VCI), as remote sensing data have better spatial and temporal coverage compared to ground data.

The outcome of this study provides for concerned bodies the meaningful and understandable information about the frequencies of hydrological extremes at various stations in Ethiopia and their association with SOI phases. This kind of information is essential for a broad group of users who are interested in monitoring, mitigation and management of droughts and floods in the country.

## Acknowledgements

I am grateful for the data I received from National Meteorology Agency (NMA) of Ethiopia. I would like to express my heartfelt thanks to all Meteorology and Hydrology department staff members at Arba Minch University who have been contributing a lot to the success of this work.

## References

Bekele F. (1997) Ethiopian Use of ENSO Information in Its Seasonal Forecasts. Internet J African Stud.

Edwards, Daniel, C., McKee and Thomas, B., (1997) Characteristics of 20th Century Drought in the United States at Multiple Time Scales. Climatology Report No. 97-2, Department of Atmospheric Science, Colorado State University, Fort Collins, CO 80523-1371.

Hayes, M., Svoboda, M., Wilhite, D. and Vanyarkho, O., (1999) Monitoring the 1996 drought using the standardized precipitation index Bulletin of the American Meteorological Society 80(3):429–438.

McKee, T.B.; N.J. Doesken; and J. Kleist. (1993) The relationship of drought frequency and duration to time scales. Preprints, 8th Conference on Applied Climatology, pp. 179–184. January 17–22, Anaheim, California

McKee, T.B.; N.J. Doesken; and J. Kleist. (1995) Drought monitoring with multiple time scales. Preprints, 9th Conference on Applied Climatology, pp. 233–236. January 15–20, Dallas, Texas.

Seiler, R., Kogan, F. and Sullivan, J., (1998) AVHRR-based vegetation and temperature Condition indices for drought detection in Argentina Remote Sensing: Inversion Problems and Natural Hazards Advances in Space Research 21(3): 481–484.

Semu, A.M. (2007) Flood Forecasting and Early Warning System (FFEWS) an Alternative Technology for Flood Management System and Damage Reduction in Ethiopia: A Concept Note, LARS 2007, 36–41.

Stone RC, Hammer GL, Marcussen T. (1996) Prediction of global rainfall probabilities using phases of the southern oscillation index. Nature, 384, 252–255.

Tannehill, I.R., (1947) Drought, its causes and effects (Vol. 64, No. 1, p. 83). LWW.

Wilhite, D. A. (1993) “Planning for drought: A methodology.” In D. A. Wilhite, ed. Drought Assessment, Management and Planning: Theory and Case Studies, pp. 87–109. Kluwer Academic Publishers, Dordrecht, the Netherlands.

Wolde-Georgis T, Aweke D, Hagos Y (2000) Reducing the Impacts of Environmental Emergencies through Early Warning and Preparedness: The Case of the 1997-98 El Niño. The Case of Ethiopia. July 10.

